# Genome-wide prediction of pathogenic gain- and loss-of-function variants from ensemble learning of a diverse feature set

**DOI:** 10.1101/2022.06.08.495288

**Authors:** David Stein, Çiğdem Sevim Bayrak, Yiming Wu, Meltem Ece Kars, Peter D. Stenson, David N. Cooper, Avner Schlessinger, Yuval Itan

## Abstract

Gain-of-function (GOF) variants give rise to increased or novel protein functions whereas loss-of-function (LOF) variants lead to diminished protein function. GOF and LOF variants can result in markedly varying phenotypes, even when occurring in the same gene. However, experimental approaches for identifying GOF and LOF are generally slow and costly, whilst currently available computational methods have not been optimized to discriminate between GOF and LOF variants. We have developed LoGoFunc, an ensemble machine learning method for predicting pathogenic GOF, pathogenic LOF, and neutral genetic variants. LoGoFunc was trained on a broad range of gene-, protein-, and variant-level features describing diverse biological characteristics, as well as network features summarizing the protein-protein interactome and structural features calculated from AlphaFold2 protein models. We analyzed GOF, LOF, and neutral variants in terms of local protein structure and function, splicing disruption, and phenotypic associations, thereby revealing previously unreported relationships between various biological phenomena and variant functional outcomes. For example, GOF and LOF variants exhibit contrasting enrichments in protein structural and functional regions, whilst LOF variants are more likely to disrupt canonical splicing as indicated by splicing-related features employed by the model. Further, by performing phenome-wide association studies (PheWAS), we identified strong associations between relevant phenotypes and high-confidence predicted GOF and LOF variants. LoGoFunc outperforms other tools trained solely to predict pathogenicity or general variant impact for the identification of pathogenic GOF and LOF variants.

## MAIN

Genetic variations exert diverse functional effects on gene products and can impact protein stability, interactions with binding partners, catalytic activity, among many other properties^1^. It is essential to investigate the functional consequences of genetic variations to understand their impact on the diverse array of observed human disease phenotypes. In particular, the functional consequences of genetic variations include two broad categories: gain-of-function (GOF) variants, characterized by enhanced or novel protein activity, and loss-of-function (LOF) variants which result in partial or complete knockdown of protein activity. GOF and LOF variants are of particular interest because they can give rise to distinct phenotypes in the same gene via contrasting molecular mechanisms^2^. For example, GOF mutations in the *STAT1* gene cause Chronic mucocutaneous candidiasis (CMC) - a susceptibility to candida infection of the skin, nails, and mucous membranes^2^. By contrast, LOF variants in *STAT1* result in Mendelian Susceptibility to Mycobacterial Disease (MSMD) - an immunodeficiency characterized by vulnerability to weakly virulent mycobacteria^2^. Given the established heterogeneity in phenotypic outcomes and their diverse modes of action, it is necessary to distinguish between GOF and LOF variants to develop a greater understanding of the genetic mechanisms of human disease, estimate individual genetic disease risk, identify candidate drug targets, and construct effective treatment regimens.

To date, effective, practical methods for distinguishing GOF and LOF variants are lacking. Experimental techniques are capable of accurately detecting GOF and LOF variants, but these methods are constrained by their significant cost and low throughput^3^. Rapid computational methods for assessing various aspects of variants such as pathogenicity or impact on protein structure/function have been developed^4–8^. Thus, for example, CADD^4^ leverages a range of functional annotations and conservation metrics to rank the relative deleteriousness of variants. PolyPhen-2^5^ and SIFT^6^ combine the physical characteristics of proteins with evolutionary features such as sequence conservation to predict whether a variant will impact protein structure or function. Tools such as REVEL^7^ and BayesDel^8^ combine the outputs of other predictors to generate a meta-score indicating variant pathogenicity. Yet, none of these tools have been designed for GOF and LOF classification.

Here we present LoGoFunc - the first effective predictor of variant functional impact - and generate predictions of functional outcomes for missense variants genome-wide. LoGoFunc is a machine learning model comprising an ensemble of LightGBM^9^ classifiers trained on pathogenic GOF and LOF variants identified in the literature. We collected 474 descriptors for use in the model including features derived from AlphaFold2^10^ (AF2) predicted protein structures, graph-based learning-derived network features representing interactions within the human protein interactome, measures of evolutionary constraint and conservation, and many others. We analyze the distributions of these features across GOF, LOF, and neutral variants, highlighting structural and functional features of proteins as well as features related to disease mechanisms such as splice disruption. Next, we assess LoGoFunc’s performance and demonstrate that LoGoFunc generates state-of-the-art predictions of GOF, LOF, and neutral variants and identifies pathogenic GOF and LOF variants more often than tools trained solely to predict pathogenicity or general variant impact. Then, we investigate which features most influence LoGoFunc’s predictions, and identify relationships between high confidence, predicted GOF and LOF variants and patient phenotypes. We provide precomputed GOF, LOF, and neutral predictions for all canonical missense variants in the human genome, which are freely available for rapid retrieval and analysis at https://itanlab.shinyapps.io/goflof/.

## RESULTS

### Labeled GOF, LOF, and neutral variant dataset curation

LoGoFunc was trained on a dataset of pathogenic GOF and LOF variants, collected from the literature via a natural language processing (NLP) pipeline^11^. In brief, the NLP pipeline parses abstracts associated with high-confidence, disease-causing variants derived from the Human Gene Mutation Database^12^ (HGMD) Professional version 2021.3, searching for terminology denoting GOF and LOF (Figure 1a). In total, 1,492 GOF and 13,524 LOF mutations were collected and labeled. In addition, 13,361 putatively neutral variants were randomly selected from the genes in which the labeled GOF and LOF variants occur, from gnomAD v2.1^13^ exome sequences (Figure 1b). We used Ensembl’s Variant Effect Predictor^14^ (VEP) to map the genomic coordinates of each variant to impacted genes and proteins where applicable and to retrieve molecular positioning information (residue position, transcript position, etc.) for each variant in the dataset. Leveraging this positional information, we further annotated each variant with 474 different features (Supplementary Table 1). These include protein structural features such as residue solvent accessibility and total residue contacts calculated from AF2^10^ predicted protein structures, gene-level features such as gene haploinsufficiency, variant-level features including splicing effects and inheritance patterns, and network features encapsulating the STRING^15^ protein-protein interaction network (Figure 1b). The annotated variants were split into label-stratified, gene-disjoint training and testing sets comprising 90% and 10% of the full dataset, respectively (Figure 1b).

**Figure 1:**
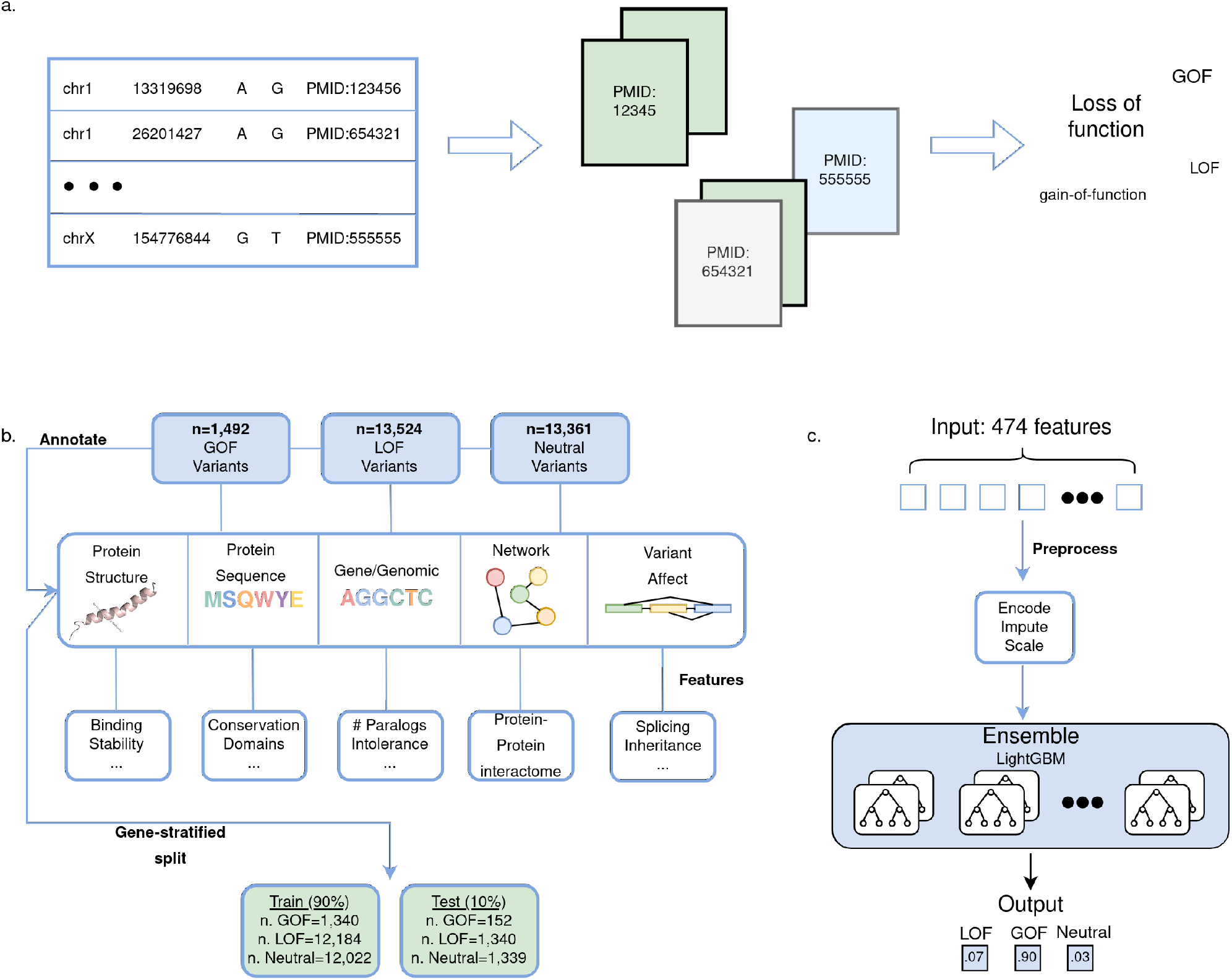
LoGoFunc workflow and model architecture. **a**. Pipeline for the collection of labeled pathogenic GOF and LOF variants. Related abstracts for high confidence pathogenic variants from the HGMD^12^ were searched for nomenclature denoting gain or loss of function. **b**. Dataset preparation and annotation. 1,492 GOF, 13,524 LOF, and 13,361 neutral variants were obtained from the GOF/LOF database^11^, HGMD^12^, and gnomAD^13^. Using VEP^14^ and other tools, variants were annotated with protein structural and functional features derived from AlphaFold2^10^ models or from sequence, with gene- and genomic-level features, variant-level features, and network-derived protein interaction features. The annotated data were split into training and test sets comprising 90% and 10% of the dataset respectively, stratified by variant label. **c**. Model architecture and output. Variants input to the model are represented as an array of the 474 collected features. These features are encoded, imputed, and scaled prior to prediction. The model consists of an ensemble of 27 LightGBM^9^ classifiers. A probability is output for each class, GOF, LOF, and neutral. Created with BioRender.com.

### GOF, LOF, and neutral variants stratified by protein features

We postulated that structural and functional features of proteins predicted or derived from protein sequences and AlphaFold2^10^ structural models may help to stratify GOF, LOF, and neutral variants. To investigate the varying impact on protein structure and function as well as potential differential localization within distinct protein regions, we examined protein features by calculating enrichments for each variant class, determined via Fisher’s exact test (Figure 2a). In total, GOF, LOF, and/or neutral variants demonstrated significant enrichments or depletions across 17 features derived from AF2^10^ predicted protein structures and across 20 protein features derived from protein sequences or otherwise describing the proteins (Figure 2a). For example, LOF variants were significantly more likely to be predicted by DDGun^16^ to have a destabilizing effect on proteins and to occur in highly conserved residues as determined by multiple sequence alignments generated by MMSeqs2^17^ (Figure 2a). GOF variants were found to be significantly more likely to occur in homomultimeric proteins and *α*-helices among other features (Figure 2a). Interestingly, both GOF and LOF variants were significantly more likely to have a high number of pathogenic HGMD^12^ variants in their spatial proximity, whereas neutral variants were significantly more likely to have a high number of gnomAD^13^ variants in their immediate vicinity. This phenomenon is exemplified by the Vasopressin V2 receptor protein in which pathogenic and putatively neutral variants can be qualitatively observed to localize to distinct regions of the 3D AlphaFold2^10^ protein structure (Figure 2b). Finally, neutral variants were significantly enriched for several features including occurrence in disordered proteins regions, and significant depletion in Pfam^18^ or InterPro^19^ domains among other features (Figure 2a). We additionally performed Fisher’s exact test with neutral variants excluded so as to compare only pathogenic GOF and LOF variants, and noted significant differences between GOF and LOF variants for seven structure-associated features and seven sequence- or otherwise associated features (Supplementary Figure 1).

**Figure 2:**
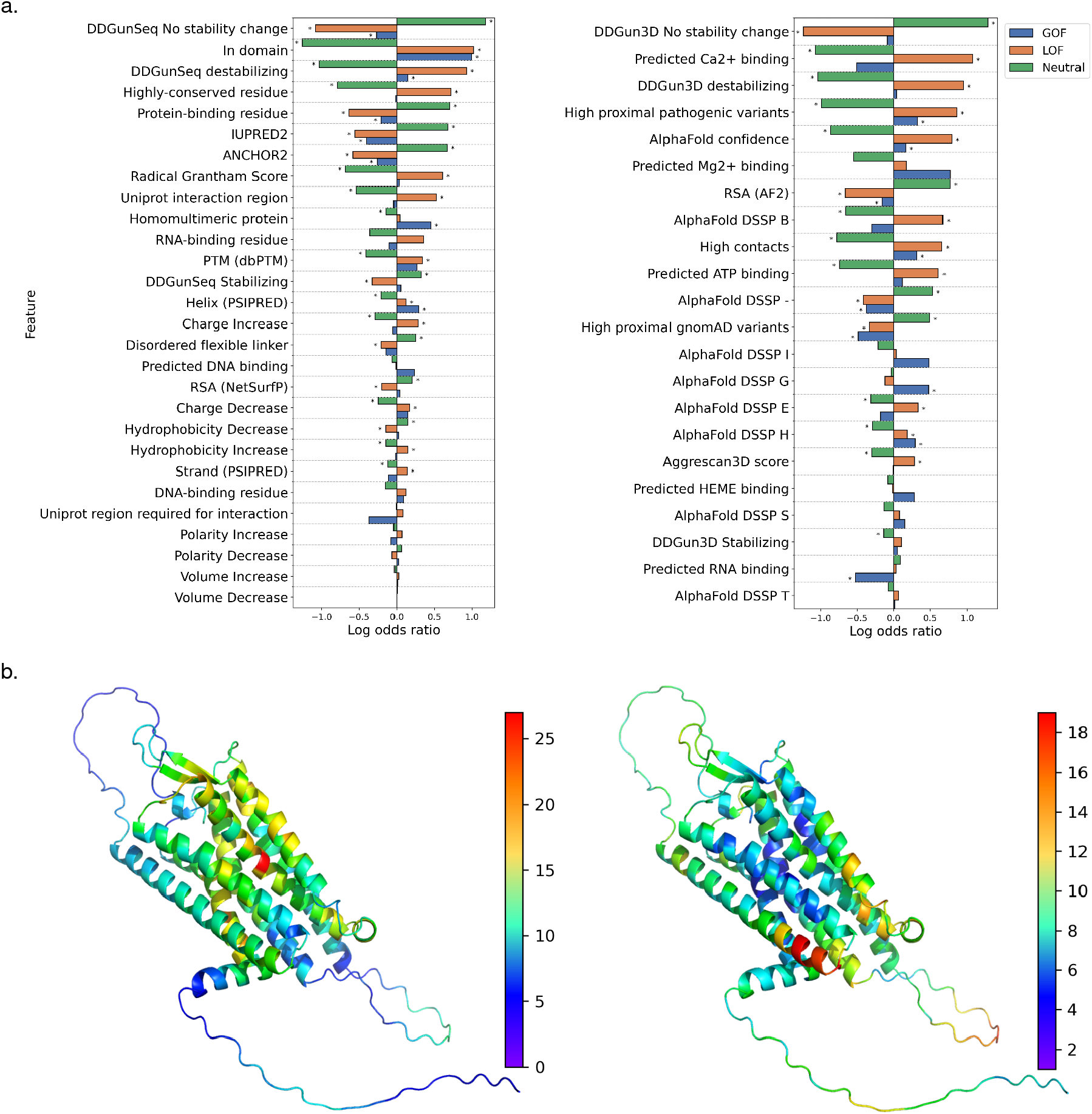
Structure- and sequence-based protein feature analysis. **a**. Enrichments and depletions for protein structural and functional features used by the LoGoFunc model. GOF (blue), LOF (orange), and neutral (green) log odds ratios are displayed for each feature. Significant enrichments and depletions are denoted by asterisks. Significance was calculated with Fisher’s exact test, Benjamini-Hochberg corrected^53^ to allow for multiple comparisons. (Left) Features derived from protein sequences or protein interaction data. (Right) Features derived from AlphaFold2 protein structures. **b**. AlphaFold2 predicted structure of the Vasopressin V2 receptor protein. (Left) Residues colored by the number of HGMD pathogenic variants occurring in the nine closest neighboring residues in space. (Right) Residues colored by the number of gnomAD variants occurring in the nine closest neighboring residues in space.

Interestingly, GOF variants were enriched and LOF variants were depleted in Pfam^18^ or InterPro^19^ domains, in *α*-helices, in homomultimer-forming proteins, and for residues not affecting protein stability based on sequence-based and structural evidence (Supplementary Figure 1). Conversely, we found that LOF variants were enriched for destabilizing amino acid substitutions, for highly conserved residues and radical Grantham^20^ PSSM substitutions, for high AF2^10^ predicted local distance difference test scoring (pLDDT) residues, and in β-strands (Supplementary Figure 1).

### GOF, LOF, and neutral variant effects on splicing

Splice-disrupting variants have been reported to constitute the second largest class of known disease-causing mutations, and have been found to yield both GOF and LOF phenotypes^21,22^. Given the importance of splice disruption as a general causal disease mechanism, we investigated the distribution of splicing-related features among the classes. Notably, LOF variants were located most closely to splice sites followed by GOF and neutral variants, respectively (p-values 2.3613E-09, 1.1287E-08) (Figure 3a, 3d). Further, LOF variants were significantly enriched for the loss of cryptic splice acceptor and donor sites (p-values 9.2849E-04, 2.5229E-07) - potentially important mechanisms of alternative splicing – and significantly depleted for the gain of cryptic splice acceptor and donor sites (p-values 2.7425E-08, 1.5519E-15) (Figure 3a). By contrast, neutral variants were enriched for the gain of splice acceptor and donor sites. The enrichment of neutral variants for the gain of cryptic splice sites (CSS) is potentially explicable in terms of the ability of canonical splice sites to suppress CSS activation^23^. Thus, these CSSs acquired via neutral mutations may not have a significant impact on transcript expression. After removing variants not predicted to impact splicing, LOF variants were predicted to lead to a greater decrease in the proportion of spliced-in (Ψ) than GOF or neutral variants based on estimates from the MMSplice^21^ exon, donor, and acceptor predictors (Figure 3c). LOF variants were similarly predicted to lead to a greater decrease in Ψ than neutral variants based on the donor-intron and acceptor-intron MMSplice^21^ predictions. GOF variants lead to a greater decrease in Ψ than neutral variants based on the exon and donor predictions (Figure 3c). These results indicate that LOF variants in particular, and to a lesser extent GOF variants, may exert their pathogenic effects via the disruption of canonical splicing patterns.

**Figure 3:**
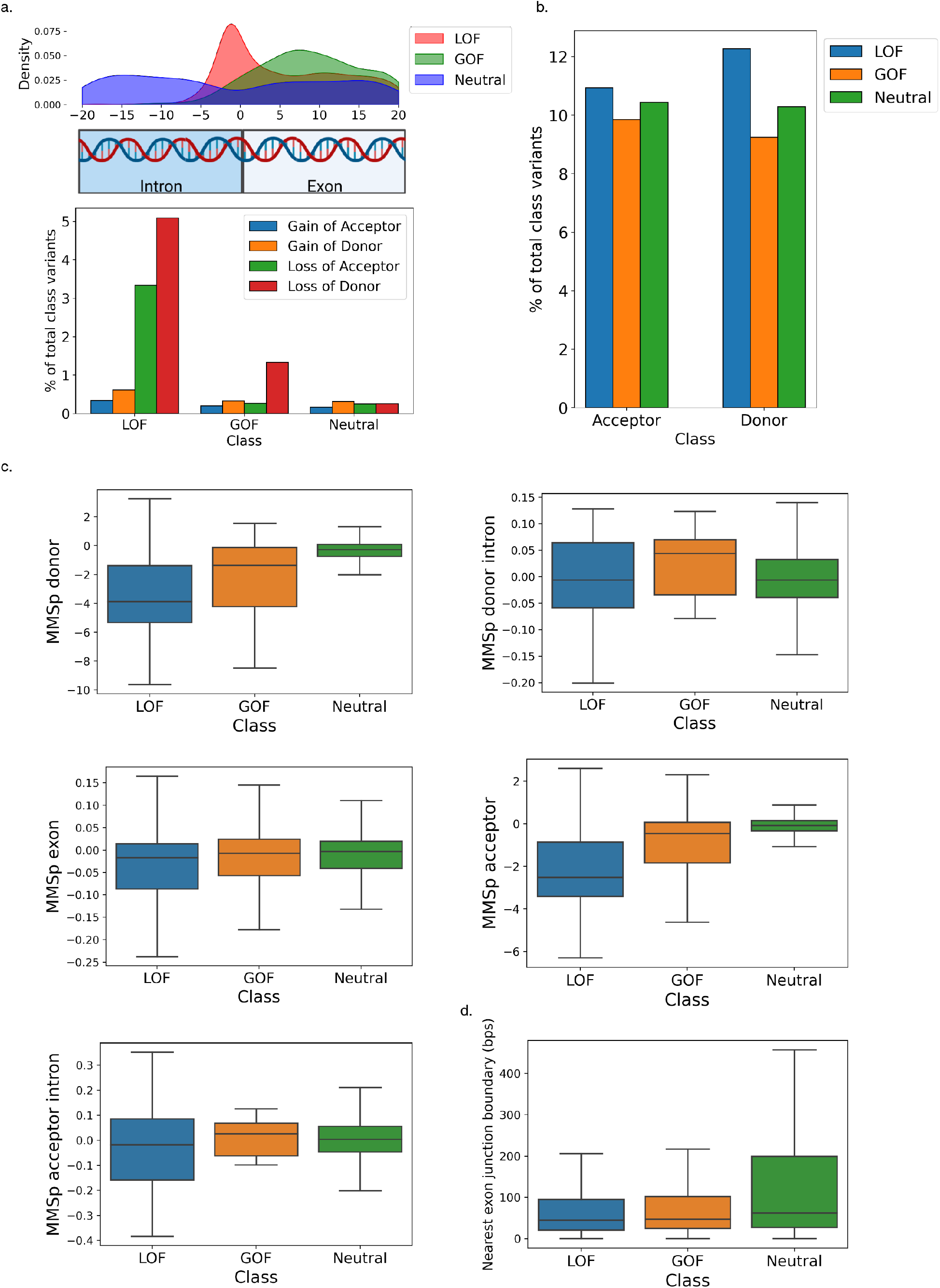
Association between variant type and impact on splicing. **a**. (Top) Density of GOF, LOF and neutral variants within 20 base-pairs of a splice junction. (Bottom) Proportion of GOF, LOF, and neutral variants predicted to yield a gain of splice acceptor or donor or a loss of splice acceptor or donor. **b**. Percentage of GOF, LOF, and Neutral variants in proximity (20 base-pairs) to acceptor and donor splice sites. **c**. MMSplice^21^ sub-model alternate minus reference logit percent-spliced-in predictions for variants predicted to impact splicing. **d**. Distance to the nearest exon junction boundary in nucleotides by variant class. Boxes denote quartiles, whilst whiskers extend to the limits of the distribution with outliers not shown when greater than 1.5 times the interquartile range from the low and high quartiles respectively. Created with BioRender.com.

### Training, architecture, and performance of LoGoFunc

The LoGoFunc model is composed of 27 LightGBM^9^ classifiers, and learned signal discriminating GOF, LOF, and neutral variants. Variants are represented as an array of 474 features that are encoded, imputed, and scaled before being input to the model which outputs three values corresponding to the predicted probability that the input variant results in a GOF, LOF, or neutral phenotype, respectively (Figure 1c).

LoGoFunc achieved notable success in classifying GOF, LOF, and neutral variants. Considering the class imbalance in the dataset, we calculated the average precision scores on the held-out testing data for each class (AP). As expected, predicting GOF variants proved to be the most challenging task as GOF variants were the least represented in the training dataset. However, LoGoFunc still performed well with AP values of .52, .93, and .96 for GOF, LOF, and neutral variants respectively (Supplementary Figure 2a). We also calculated the F1-score and Matthew’s Correlation Coefficient for LoGoFunc’s predictions of variants from each class. LoGoFunc realized F1-scores of .56, .87, and .89 and Matthew’s Correlation Coefficients of .54, .75, and .80 for GOF, LOF, and neutral variants respectively. To aid in the interpretation of LoGoFunc’s predictions, we calculated 95% confidence intervals for determining cutoffs for each class, as well as 95% confidence intervals for determining GOF, LOF, and neutral prediction cutoffs per gene (Supplementary Table 2).

### Benchmark against variant assessment algorithms

Currently, there are no high-throughput computational predictors trained to classify pathogenic GOF and LOF variants^11^. We therefore compared LoGoFunc to ten established predictors of pathogenicity/deleteriousness: CADD^4^, SIFT^6^, PolyPhen2^5^, DANN^24^, BayesDel^8^, ClinPred^25^, GenoCanyon^26^, MetaSVM^27^, PrimateAI^28^, and REVEL^7^. To equitably assess each method’s ability to discriminate GOF and LOF, we selected the subset of 1,092 GOF, LOF, and neutral variants from the test set for which all predictors provided a score. Of these variants, 136 were GOF, 545 were LOF, and 411 were neutral. Importantly, these variants were all missense, as the majority of compared methods provide predictions only for missense variants. We tested each method’s performance in separating LOF from neutral, GOF from neutral, GOF from LOF variants, and both GOF and LOF combined from neutral. LoGoFunc achieved an AP of .87 for LOF *vs*. neutral (Figure 4a) and .82 for GOF *vs*. neutral (Figure 4b). The next best tool, REVEL^7^, achieved AP values of .87 and .55 for LOF and GOF *vs*. neutral respectively (Figure 4a,b). We also compared the model’s ability to separate GOF from LOF variants. LoGoFunc achieved an AP of .63 followed by GenoCanyon^26^ with a score of .25 (Figure 4c). We calculated the one-*vs*.-all AP for the neutral variants against the GOF and LOF variants. Once again, LoGoFunc scored highest with an AP of .91, followed by REVEL^7^ with an AP of .88 (Figure 4d). LoGoFunc performed as well as or outperformed the other models for each comparison, particularly for the separation of GOF variants from neutral variants and GOF and LOF variants from each other, indicating that training on labeled GOF and LOF variants may yield a model better suited for identifying protein gain- and loss-of-function as a result of genetic variation.

**Figure 4:**
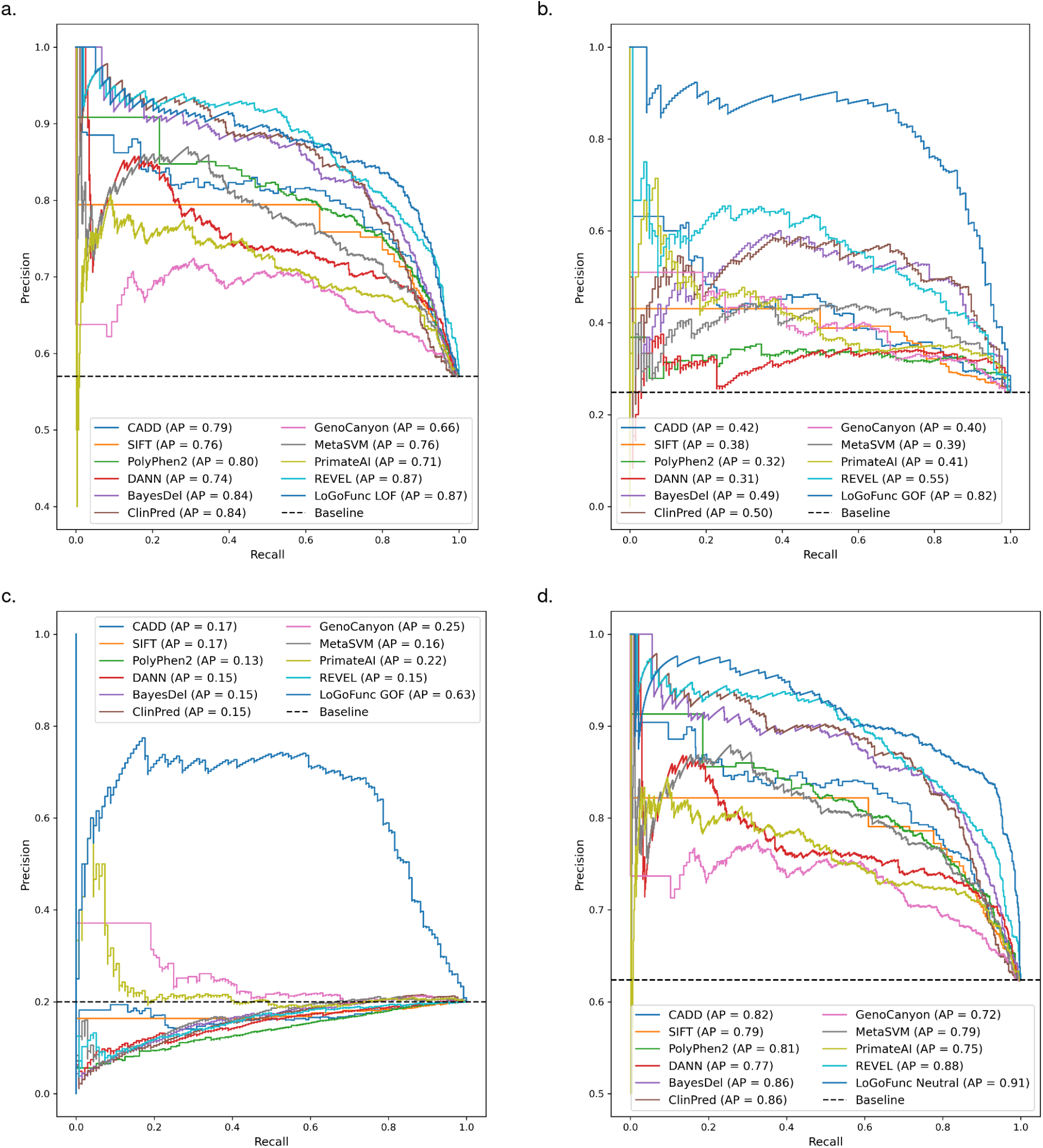
Benchmarking LoGoFunc. Precision-recall curves comparing the discriminatory power of various pathogenicity prediction methods and LoGoFunc on a set of variants from the test set for which predictions were available from all compared tools. **a**. LOF (n. 545) *vs*. neutral (n. 411). **b**. GOF (n. 136) *vs*. neutral (n. 411). **c**. GOF (n. 136) *vs*. LOF (n. 545). **d**. GOF (n. 136) and LOF (n. 545) combined *vs*. neutral (n. 411).

### LoGoFunc leverages diverse biological signals for prediction

To gain further insight into the model’s performance, we estimate the impact of each included feature on LoGoFunc’s predictions with SHAP^29^ – a game theoretic approach for the derivation of explanations for machine learning models (Figure 5a). We observed that LoGoFunc learned from a diverse array of features describing the genes and proteins containing variants, and the variant impact upon these elements. These included functional, conservation, structural, and systems-based/network features, among others (Supplementary Table 3, Figure 5a). For example, the top feature across classes was the consequence score collected from the CADD^4^ database of variant annotations which describes the severity of a variant according to sequence ontology^30^ consequence terms (Figure 5a). Other important variant features include predictions indicating pathogenicity from CADD^4^, VEST4^31^, M-CAP^32^, and MVP^33^, the MOI-pred^34^ mode of inheritance prediction of variants underlying autosomal dominant (AD) and autosomal recessive (AR) disease, and various measures of conservation from tools such as GERP^35^, PhyloP^36^, and PhastCons^36^ (Figure 5a). Several gene-level features were important for the model including the number of gene paralogs, the *de novo* excess rate^37^, the mutation significance cutoff^38^ 95% confidence interval, and the indispensability score^39^ – all of which have previously been implicated in the stratification of pathogenic GOF and LOF variants and neutral variants^11^ (Figure 5a). In addition, LoGoFunc’s predictions were influenced by features indicating variant effects on protein structure and function such as the predicted variant impact on protein stability, the number of HGMD^12^ pathogenic or gnomAD^13^ variants proximal to variant impacted residues in 3D space, AlphaFold2^10^ pLDDT scores which indicate AF2’s^10^ per-residue prediction confidence, and overlapping Pfam^18^ or InterPro^19^ domains (Figure 5a). Notably, protein-protein interaction (PPI) network features also had a significant impact on the model. We processed the STRING^15^ protein-protein interaction (PPI) network using node2vec^40^ resulting in 64 tabular features summarizing the human protein interactome weighted by the probability of interaction between each pair of putatively interacting proteins. Several dimensions of the transformed PPI network appeared in the list of top features as determined by SHAP^29^ (Figure 5a).

**Figure 5:**
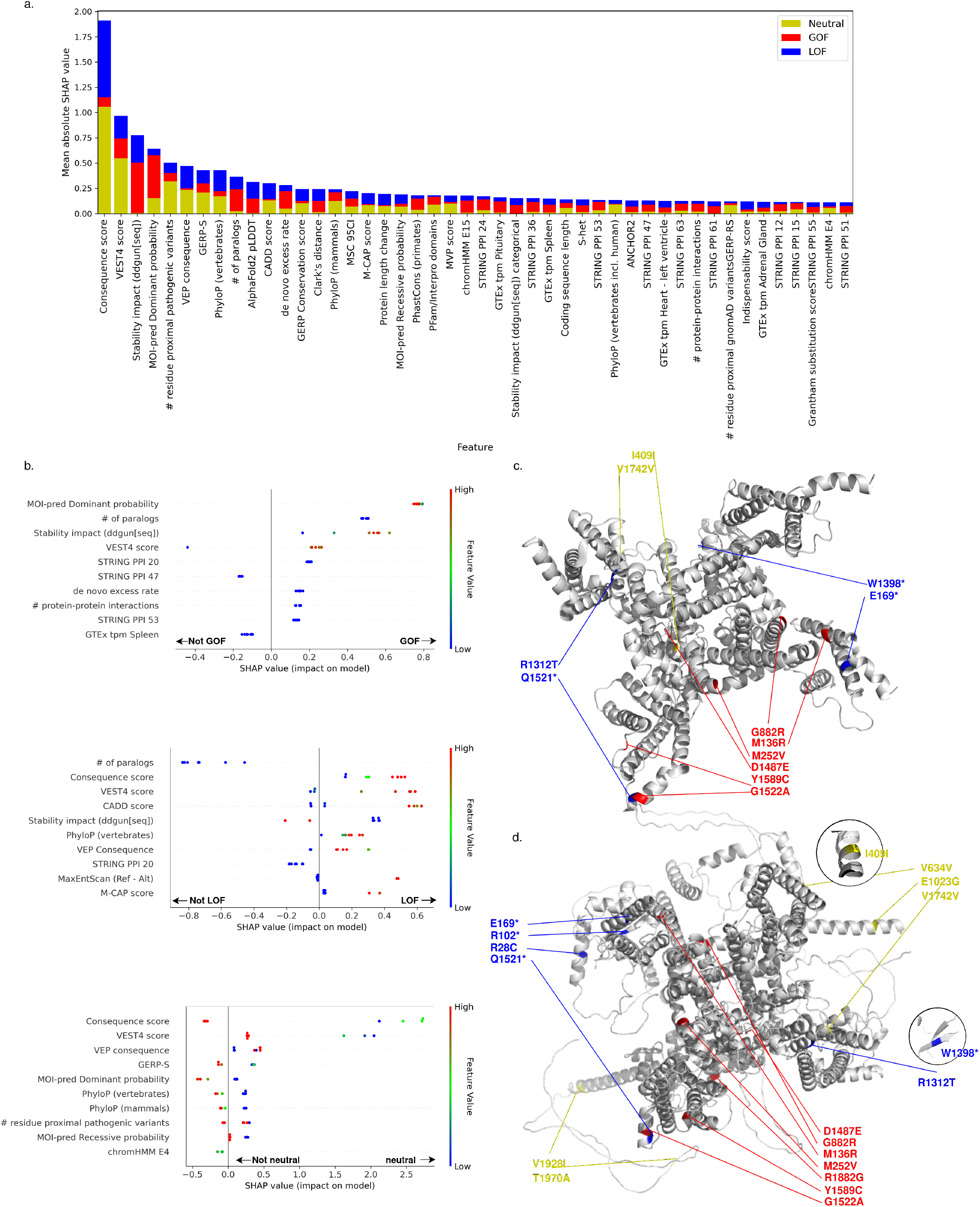
Explanation of LoGoFunc predictions. **a**. SHAP values by class for features with combined SHAP values in the 90^th^ percentile and above. **b**. (Top) The SHAP values for the top ten features for the seven GOF variants found in the ion channel SCN2A in the test set. (Middle) The SHAP values for the top ten features for the eight LOF SCN2A variants in the test set. (Bottom) The SHAP values for the top ten features for the seven neutral SCN2A variants in the test set. **c**. The experimentally determined structure of SCN2A^43^ with the represented GOF (red), LOF (blue), and neutral (yellow) SCN2A variants from the test set. **d**. The SCN2A model from the AlphaFold2 prediction database annotated with the represented GOF (red), LOF (blue), and neutral (yellow) SCN2A variants from the test set.

To further investigate the model’s predictions within genes, we examined the 22 variants included in our test set from sodium voltage-gated channel alpha subunit 2 (SCN2A) - an important transmembrane protein implicated in seizure disorders^41^ and autism spectrum disorders^42^. Of these 22 variants, VEP^14^ indicated twelve to be missense, four to be stop-gains, two to be splice donor site variants, three to be synonymous, and one to be intronic. Twelve of the coding variant positions are included in the experimentally determined structure (PDB identifier 6J8E^43^) (Figure 5c). Because the other ten variants are located in regions not covered by the structure, we analyzed the structural model generated by AF2^10^ (Figure 5d), which includes the full-length protein. Remarkably, LoGoFunc successfully classified all seven SCN2A pathogenic GOF variants, all seven SCN2A neutral variants, and six of eight pathogenic LOF variants, misclassifying two LOF variants as GOF. We then examined the top ten features indicated by SHAP^29^ to be contributing to the model’s predictions for the GOF, LOF, and neutral variants separately (Figure 5b). Again, we found a mixture of gene, protein, variant, and network features influenced the model’s predictions. Specifically, a range of MOI-pred^34^ mode of inheritance prediction of variants pathogenic for AD inheritance, mid-range and higher DDGun^16^ predictions indicating less protein destabilization, and high VEST4^31^ scores among others influenced the model to predict the SCN2A GOF variants as GOF. Similarly, several features prompted the model to predict the LOF variants to be LOF, including high consequence scores indicating higher impact on transcripts and downstream products, high VEST4^31^ and CADD^4^ scores, low DDGun^16^ scores indicating a greater destabilizing effect on proteins, and high vertebrate PhyloP^36^ scores indicating higher conservation. Notably, high MaxEntScan^44^ difference scores contributed to the model’s LOF predictions, consistent with VEP’s characterization of two of the LOF variants as splice donor site variants. The model’s predictions were most influenced towards neutrality by lower consequence scores, lower VEST4^31^ scores, lower GERP-S^45^, and vertebrate and mammalian PhyloP^36^ scores, and lower MOI-pred^34^ scores among other features.

### PheWAS corroborates LoGoFunc predictions

We performed phenome-wide association study (PheWAS) analyses on a subset of predicted GOF and LOF missense variants which were either absent from, or indicated as variants of uncertain significance (VUS) in, ClinVar^46^ (Supplementary Table 4). In brief, PheWAS evaluates the association between a genetic variant and a set of phenotypes. Although our analysis was insufficiently powered for phenome-wide significance, as expected due to the low frequency of our variants, we nevertheless uncovered meaningful associations between our variants and relevant phenotypes (Figure 5a). For example, we observed that the predicted LOF variant c.1648G>A (rs563131364) in the *SLC12A3* gene, that encodes a sodium-chloride co-transporter, is strongly associated with increased risk for several phenotypes including severe chronic kidney disease (p=0.001, LO=1.723, ICD=N184), abnormal blood chemistry (p=0.003, LO=1.189, ICD=R7989), and retinal edema (p=0.006, LO=2.377, ICD=H3581), among several other conditions. Conversely, the mutation was found to be protective with respect to pure hypercholesterolemia (p=.0396, LO=-1.512, ICD=E7800). The c.1648G>A variant has a CADD^4^ PHRED score of 29.3 and an MSC^38^ 95 CI of 13.89, indicating that the variant may be pathogenic taking into consideration the genic context of *SLC12A3* (Figure 5b). c.7471C>T (rs201746476) is a predicted GOF variant in the *PIEZO1* gene, which encodes a mechanosensitive ion channel and has previously been linked to arrhythmia when overexpressed^47^. c.7471C>T was associated with risk for palpitations (p=0.009, LO=2.912, ICD=R002), abdominal pain (p=0.028, LO=1.842, ICD=R109), and viral hepatitis C susceptibility (p=0.039, 0.043, LO=2.438, 2.389, ICD=B182, B1920). Notably, GOF mutations in *PIEZO1* have been shown to impair hepatic iron metabolism^48^, a mechanism of viral hepatitis C infection^49^. Similar to c.1648G>A, c.7471C>T had a CADD PHRED score of 25.4, well over the MSC^38^ 95 CI of 2.185 for the *PIEZO1* gene.

## DISCUSSION

Describing the functional consequences of genetic variations is critical for the development of a better understanding of disease mechanisms. We have developed LoGoFunc, a rapid and accurate predictor of GOF, LOF, and neutral variants, and used it to analyze various features associated with these variant types. Four key findings emerge from this work.

First, we observe that pathogenic GOF, LOF, and neutral variants inhabit varying structural and functional regions of proteins, exert differing effects on protein structure, and inhabit proteins with different protein-protein interaction characteristics (Figure 2, Supplementary Figure 1). Specifically, LOF variants consistently demonstrate a greater propensity for the disruption of protein structure and/or function. Particularly, as predicted by DDGun^16^ leveraging both sequence-based and structural evidence, LOF variants are significantly more likely to have a destabilizing effect on protein structure and significantly less likely to stabilize or result in a negligible effect on protein structure. LOF variants are enriched for highly conserved residues as identified by multiple sequence alignments from MMSeqs2^17^ and for more radical amino acid substitutions as calculated from the Grantham^20^ position-specific scoring matrix. Similarly, LOF variants are enriched for known post-translationally modified residues (PTMs) and are more likely to be buried in protein structures as evidenced by residue relative solvent accessibility (RSA) predictions from NetSurfP^50^ and RSA calculated by DSSP^51^ using AF2^10^ structures. GOF variants compared to LOF, while enriched in potentially functionally important Pfam^18^ domains, appear to impact protein structure less radically. Indeed, compared to LOF variants, GOF variants were depleted for predicted protein destabilizing substitutions, highly conserved residues based on MSAs, and radical Grantham^20^ substitutions. Interestingly, when considering both sequence-based predictions and evidence derived from AF2^10^ structures, we found GOF variants to be enriched in *α*-helices, and LOF variants to be enriched in β-strands. Previous studies have demonstrated mutations in *α*-helices to be less structurally impactful than mutations occurring in β-strands^52^, consistent with the characterization of GOF and LOF variants established by other features.GOF variants were also enriched in proteins capable of forming homomultimers suggesting a potential dominant negative pattern of gain of function for some of the variants and further emphasizing the necessity to investigate protein interactions when assessing variant functional impact. Together, these observations indicate significant divergence between GOF and LOF variants in their mode of pathogenicity at the protein level and suggest several mechanisms that may guide and inform the investigation of individual variants. Further, these results demonstrate that AF2^10^ predicted protein structures may provide significant biological signal in variant assessment tasks and can facilitate the extraction of protein structural features proteome-wide.

Second, LoGoFunc demonstrates strong performance on an independent test set of GOF, LOF, and neutral variants and, by considering functional outcomes during training, is better able to predict the functional impact of genetic variants than tools trained under a binary benign/pathogenic paradigm (Figure 4). Interestingly, the benchmarked tools in our analysis performed better on LOF variants than GOF. It has previously been demonstrated that several pathogenicity predictors such as CADD^4^ and REVEL^7^ tend to predict LOF variants as pathogenic or deleterious more often than GOF variants, whereas GOF variants are more often predicted to be benign^11^. This may be due in part to the underrepresentation of GOF variants in the training data used by these tools where applicable or may arise because GOF variants may be difficult to separate from neutral variants using the features or methods employed by these tools. Importantly, these results suggest that LoGoFunc may be particularly useful for predicting GOF variants, as it may be capable of identifying pathogenic GOF variants that other pathogenicity predictors would tend to misclassify.

Third, our analysis identified previously undocumented associations between various biological features and the functional outcomes of genetic variants. We assessed the importance of the features used to train LoGoFunc and found that the model learns from a diverse array of gene-, protein-, and variant-level features including functional, conservation, structural, and network information (Figure 5). For example, we processed the STRING^15^ protein-protein interaction (PPI) network using node2vec^40^ to summarize the human protein interactome. Whereas some models have included binary indications of the involvement of a protein in any protein interaction^53^, to our knowledge, such PPI network features are rarely used in popular pathogenicity prediction methods. Yet, many dimensions of the output are highly impactful for the LoGoFunc model, suggesting protein function at the pathway- and/or systems-level may have a bearing on variant pathogenicity and functional effect.

Concordantly, PPI features are accompanied by several other protein sequence- and structure-based features from which the model also learns, including top features such as DDGun^16^ stability impact predictions, residue proximal pathogenic variants, and the AF2 structure pLDDT values which have been shown to correlate significantly with protein structural disorder^10^. Genic context also has a substantial impact on the model’s output as evidenced by the inclusion of several gene-level features such as the gene damage index^54^, and the number of gene paralogs. Other important features, such as the per variant predictions of pathogenicity for autosomal dominant or recessive disease, align with previous characterizations of GOF and LOF variants, thereby supporting the biological plausibility of LoGoFunc’s predictions and lending credence to the novel associations we identified between various features employed by the model and GOF, LOF, and neutral variants.

Finally, we illustrate LoGoFunc’s potential utility for characterizing variants of uncertain significance (VUS) and uncharacterized variants, a major challenge in human genomics. We performed PheWAS on predicted GOF and LOF variants, which were either marked as VUS in or were absent from ClinVar^46^, using patient records from the Mount Sinai BioMe Biobank (Figure 6). We uncovered strong associations between the tested variants and relevant phenotypes. For example, the predicted LOF variant c.1648G>A (rs563131364) in *SLC12A3* was associated with severe chronic kidney disease and abnormal blood chemistry among other phenotypes. Notably, over 140 putative LOF *SLC12A3* variants have previously been identified in patients with Gitelman syndrome^55^, a disorder characterized by impaired salt reabsorption in the kidneys, including four neighboring variants in the same transmembrane helical region. Analysis of specific features also suggested that c.1648G>A may be a pathogenic, loss-of-function variant. For example, the variant has a CADD^4^ PHRED score of 29.3 and an MSC^38^ 95 CI of 13.89, indicating that the variant may be pathogenic taking into consideration the genic context of *SLC12A3*. Similarly, the variant is predicted to manifest autosomal recessive inheritance, consistent with the inheritance pattern of Gitelman syndrome^55^. Together, these results provide preliminary evidence that LoGoFunc may provide utility in the assessment of VUS and uncharacterized variants in addition to providing predictions of functional effect.

**Figure 6:**
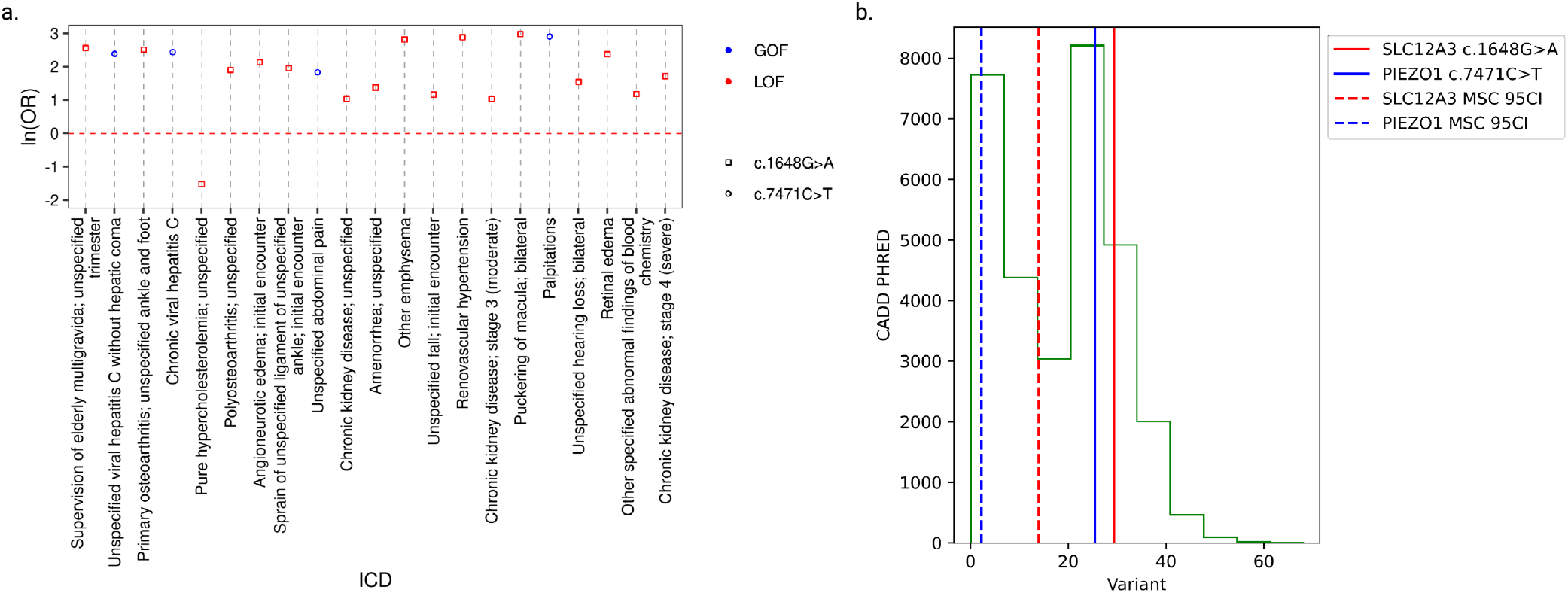
Relationship between variant type and phenotypes. **a**. Associations between high- confidence, predicted GOF (c.7471C>T) and LOF (c.1648G>A) variants and phenotypes as determined by PheWAS analysis of patients in the BioMe biobank. **b**. Distribution of CADD^4^ PHRED scores in the dataset (green). CADD^4^ PHRED scores and MSC^38^ 95% CI cutoffs for c.7471C>T (solid and dashed blue lines) and c.1648G>A (solid and dashed red lines).

In summary, we have developed LoGoFunc, a predictor of GOF, LOF, and neutral variants. Our model performs favorably compared to commonly used computational tools designed for the assessment of genetic variation and demonstrates strong predictive power across metrics on our test set of GOF, LOF, and neutral variants. We assessed the contribution of various features to the model’s output and found that LoGoFunc learns from a diverse array of structural, functional, sequence-based, and systems-level information, indicating that these features have a bearing on the functional outcome of genetic variants. Further, we demonstrated significant localization of GOF, LOF, and neutral variants in 3D structural and functional sites in proteins, and demonstrated LoGoFunc’s ability to assess previously uncharacterized variants. Our findings corroborated previously reported molecular mechanisms resulting in the gain or loss of function and also suggest novel mechanisms that may shed light on disease etiology. We applied our method to 82,468,698 canonical missense mutations in the human genome, and provide our predictions at https://itanlab.shinyapps.io/goflof/.

## METHODS

### Dataset assembly

We obtained 11,370 labeled pathogenic GOF and LOF variants from Bayrak et al^11^. To supplement this dataset, we collated the 65,075 variants that were deposited in the HGMD^12^ Professional version 2021.3 database specifically in 2020 and 2021 and assigned labels using the same strategy that Bayrak et al^11^ employed. From these variants, we first selected 32,911 disease-causing class (DM) variants. We then used the Spacy 3.0.6 natural language processing (NLP) library to search for GOF- and LOF-related nomenclature in associated publications for each DM variant. Using the phrase-based matching algorithm PhraseMatcher, we iteratively searched the paper titles and abstracts from all associated publications for the patterns “gain of function(s)”, “gain-of-function(s)”, “GOF”, “loss of function(s)”, “loss-of-function(s)”, and “LOF” with text converted to lowercase to allow for case sensitivity. When at least one of the publications indicated GOF or LOF, we labeled the corresponding variant accordingly. When there was a disagreement, i.e. a variant was found as GOF in one abstract and LOF in another abstract, the variant was excluded from the dataset.

Putatively neutral variants were selected from the gnomAD v2.1^13^ exome sequencing data. gnomAD^13^ variants were selected from genes represented by the labeled GOF and LOF variants after filtering HGMD^12^ pathogenic variants from the gnomAD^13^ dataset. A minimum of two gnomAD^13^ variants and up to the number of GOF or LOF variants, whichever was the lower, were selected from each gene represented by the labeled GOF and LOF variants for a total of 13,361 putatively neutral variants. The complete labeled dataset comprising 1,492 GOF, 13,524 LOF, and 13,361 neutral variants was split into training and testing sets such that the ratio of GOF to LOF to neutral variants in the training and testing sets reflected the ratio in the complete dataset, and such that there was no overlap of represented genes between the training and testing sets. The training set and testing sets comprise 90% and 10% of the complete dataset, respectively.

### Variant annotations

Ensembl’s VEP^14^ version 106 was employed to annotate all variants according to their GRCh38 genomic coordinates. VEP^14^ provided affected transcripts, genes, and proteins, and the position of variants within these elements where applicable. VEP^14^ plugins provided pathogenicity predictions from CADD^4^, SIFT^6^, PolyPhen2^5^, and CONDEL^56^. Additional pathogenicity predictions were collected using the VEP^14^ dbNSFP^57^ plugin version 4.1a, along with variant allele frequencies, and conservation scores from PhastCons^36^, PhyloP^36^, SiPhy^58^, and GERP++^45^. VEP^14^ plugins were also used to retrieve BLOSUM62^59^ scores, GERP^35^ scores, distances from variants to the nearest exon junction boundary and the nearest transcription start site, MaxEntScan^44^ predictions, dbscSNV^60^ splice variants, and predictions of variants allowing for transcript escape from nonsense-mediated decay. AlphaFold2^10^ (AF2) structural models were downloaded from https://ftp.ebi.ac.uk/pub/databases/alphafold/latest/UP000005640_9606_HUMAN_v3.tar^61^.

The Biopython PDB module was used to load PDB^62^ formatted AF2^10^ models and to calculate various geometric properties of proteins and residues. Specifically, residue contacts were inferred when the *α*-carbons of a given pair of residues resided within 12 Angstroms of each other in 3D space. Similarly, the distance of each residue from the protein center of mass was defined as the 3D distance in Angstroms from the residue’s *α*-carbon to the protein center of mass as calculated by the Biopython PDB module. To calculate the number of proximal HGMD^12^ pathogenic and gnomAD^13^ variants in a residues 3D environment, we first mapped protein coordinates to genomic positions for the 18,901 canonical human proteins for which UniProt^63^ provides such a mapping. The number of pathogenic or gnomAD^13^ variants occurring in the nine closest residues in 3D space based on the structural models was then summed for each residue in each protein. The Biopython PDB and DSSP^51^ modules were used to extract secondary structure characterizations and relative solvent accessibility for the model residues. Putative protein-ligand binding sites were predicted using ConCavity^64^ v0.1 with the protein structural models as input (default parameters). DDGun^16^ and GraphBind^65^ were similarly employed to predict variant impacts on protein stability and ligand binding residues respectively using the default parameters and the structural models. All other features were collected from their respective web servers or calculated via standalone tools (Supplementary Methods, Supplementary Table 1).

### Feature analysis and feature importance

Feature enrichments were calculated via Fisher’s exact test. Continuous features obtained from the DescribePROT^66^ database were categorized according to the cutoffs derived from proteome-wide metrics described in Zhao et. al^66^. Residues were classified as buried if their RSA was less than 20%; otherwise, they were regarded as exposed. Grantham^20^ scores for amino acid substitutions were considered to be conservative if lower than 100 and radical if greater than or equal to 100. The numbers of residue contacts were binned into categories “high” and “low” based on the median number of residue contacts across the 20,504 proteins included in the AF2^10^ *Homo sapiens* reference proteome dataset. Similarly, the number of residue proximal pathogenic variants from HGMD^12^ and residue proximal gnomAD^13^ variants were categorized as “high” or “low” based on the median value of each of these features across the 18,901 proteins for which UniProt^63^ provided a mapping between genomic coordinates and residue position. Other continuous features were categorized by assigning a cutoff according to the value recommended by the authors of the tools from which the features were derived. When no such cutoff was reported, a cutoff of 0.5 was selected for probabilistic features. Distance from exon-intron junction boundaries and MMSplice^21^ predictions were compared via one-sided two-sample T-tests. The Benjamini-Hochberg correction^67^ was applied at an alpha level of 0.05 to control for false positives as a result of multiple testing. Feature importance was assessed via the SHAP^29^ Python package version 0.41.0. Specifically, the mean SHAP^29^ values across the ensembled LightGBM^9^ models were generated via the SHAP^29^ tree explainer model.

### Preprocessing of input data

Preprocessing steps were applied to prepare sample variants for prediction. An ordinal encoder was fitted to the categorical features in the training set and used to encode the categorical features in the training and test sets. Missing values were imputed either with a constant (−1) or with the median value of the feature in the training set. Zero variance features in the training set were dropped from both the training and test sets. Finally, random oversampling was performed on the GOF and neutral variants to bring their total count in the training set equal to the majority class, LOF.

### Model selection

We performed 5-fold outer, 5-fold inner, nested cross-validation in which folds did not contain variants from the same sets of genes on the training dataset to assess the variance associated with our preprocessing pipeline, model hyperparameters, and model architecture (Supplementary Figure 3). Specifically, we evaluated the performance and generalizability of four models: RandomForest^68^, LightGBM^9^, XGBoost^69^, and Neural Networks. For each algorithm, the data preprocessing procedure and relevant hyperparameters were tuned for 200 rounds in each iteration of the inner cross-validation loop with the Optuna^70^ optimization library to maximize the macro-averaged F1-score (F1) (for hyperparameter search spaces see Supplementary Information). The F1-score is a function of the precision and recall, defined as follows, where *y* is the set of predicted sample, label pairs, and *y’* is the set of true sample, label pairs:

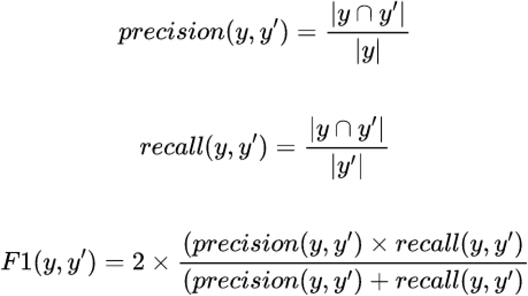

To extend the F1-score to multiclass classification, we calculated the macro-averaged F1-score, defined as follows where *L* is the set of labels:

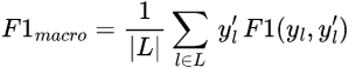

The preprocessing pipeline and hyperparameters which performed best for each model in the inner cross-validation iteration were then used to assess each model on the held-out set of the outer cross-validation loop. After all rounds of outer and inner cross-validation, the median Matthew’s correlation coefficient (MCC) and F1 were compared to determine which model performed best for the dataset. The MCC is defined as follows where *k* is the number of classes and *kl* refers to an element of the confusion matrix:

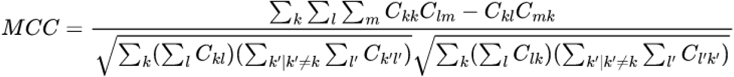

LightGBM^9^ obtained the best MCC and F1 scores across outer folds (Supplementary Figure 4). We subsequently performed the same nested cross-validation procedure described above with ensembles of 5 to 31 LightGBM^9^ models with individual model hyperparameters and the number of ensemble estimators tuned simultaneously. The ensembled LightGBM^9^ models achieved the highest MCC and F1 scores across outer folds and were selected as the final model. Subsequently, we performed the inner cross-validation procedure with all of the training data to determine the final number of ensemble estimators and model hyperparameters.

### LoGoFunc performance

LoGoFunc’s performance was assessed via average precision (AP), F1-score, and Matthew’s correlation coefficient calculated using scikit-learn version 1.1.1. AP is defined as follows, where *n* is the *n*_th_ threshold:

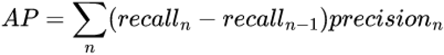

For each class, we computed these metrics as one *vs*. rest tasks where the class in question was relabeled as one and the other classes were relabeled as zero.

### Gene 95% confidence intervals

For each variant class, GOF, LOF, and neutral, we selected predictions from the training and testing sets for variants of that class. We applied the Kolmogorov-Smirnov^71^ test for goodness of fit to predictions for these variants with continuous distributions implemented in scipy^72^ version 1.0.1. For each distribution, we first estimated the distribution parameters that best modeled the predictions using scipy^72^, and then selected the parametrized distribution with the highest p-value from the Kolmogorov-Smirnov test^71^. 95% confidence intervals were then calculated using the best fitting, parameterized distribution for predictions from each class respectively and clipped between zero and one where applicable. When five or more variants were available from a given class for a given gene, we repeated the above process to calculate gene-specific 95% confidence intervals. When fewer than five variants were available for a class in a gene, we defaulted to the 95% confidence intervals calculated for predictions from the entire dataset.

### Method comparison

LoGoFunc was compared to other computational methods by assessing the AP. All GOF (n. 136) and LOF (n. 545) variants from the test set for which all compared tools provided a prediction were collected. APs were calculated, treating GOF as the positive class. To assess the performance separating neutral variants from GOF and LOF, we added all neutral (n. 411) variants from the test set for which each tool provided a prediction. APs were again calculated, this time with GOF and LOF variants as the positive classes respectively, and neutral as the negative class. Finally, we calculated the one-*vs*.-all APs with GOF and LOF variants as the positive class and neutral variants as the negative class. Most of the compared tools provide predictions in which higher scores correspond to a greater likelihood that a given variant will be damaging. However, SIFT outputs predictions between zero and one in which lower scores correspond to a greater likelihood of a damaging effect.

LoGoFunc’s neutral prediction is a value between zero and one, where higher scores indicate a greater likelihood of neutrality. Thus, to ensure consistency between all compared tools when treating neutral as the negative class and GOF and LOF as the positive class, SIFT and LoGoFunc neutral predictions were transformed by subtracting each prediction from one before assessing AP.

### PheWAS of predicted GOF and LOF variants

We analyzed 1,650 phenotypes for which there were at least 20 cases in the Mount Sinai BioMe BioBank. For each phenotype, controls were randomly sampled from non-cases to fix the ratio of cases to controls at 1:5. Overlapping individuals, i.e. those sharing phenotypes other than the phenotype of interest, were removed from the control set. We implemented principal component analysis (PCA) on 3,800 whole-exome sequencing samples in BioMe using independent variants before the PheWAS analysis and used the first five components to adjust for potential population stratification in both cases and controls. We reduced linkage disequilibrium (LD) between markers by removing all markers with r^2^>0.2 (window size 50, step size 5), as well as markers in known high LD regions. Furthermore, we retained variants with minor allele frequency (MAF) greater than 0.02 and genotyping rate greater than 95% across the dataset (excluding A/T, C/G mutations). PCA was conducted using Plink 1.9^73^. PheWAS was conducted using the R “PheWAS” package^74^.

## ACKNOWLEDGMENTS

We thank Bruce Gelb, Vikas Pejaver, Laura Huckins, Josh Milner, Nicole Zatorski, and Keino Hutchinson for their thoughtful discussion and support for this project.

## AUTHOR CONTRIBUTIONS

DS, CB, YI, and AS conceived of this project. CB and DS collected the labeled data for training and testing the model. YW and MEK carried out the PheWAS analysis. DS led the development of the model and the preparation of the manuscript. PDS and DNC provided data for training and testing. YI and AS oversaw this project and provided guidance. All authors contributed to the writing of the manuscript and approved the final manuscript.

## FIGURE LEGENDS

**Figure 1: LoGoFunc workflow and model architecture. a**. Pipeline for the collection of labeled pathogenic GOF and LOF variants. Related abstracts for high confidence pathogenic variants from the HGMD^12^ were searched for nomenclature denoting gain or loss of function. **b**. Dataset preparation and annotation. 1,492 GOF, 13,524 LOF, and 13,361 neutral variants were obtained from the GOF/LOF database^11^, HGMD^12^, and gnomAD^13^. Using VEP^14^ and other tools, variants were annotated with protein structural and functional features derived from AlphaFold2^10^ models or from sequence, with gene- and genomic-level features, variant-level features, and network-derived protein interaction features. The annotated data were split into training and test sets comprising 90% and 10% of the dataset respectively, stratified by variant label. **c**. Model architecture and output. Variants are input to the model represented as an array of the 474 collected features. These features are encoded, imputed, and scaled prior to prediction. The model consists of an ensemble of 27 LightGBM^9^ classifiers. A probability is output for each class, GOF, LOF, and neutral. Created with BioRender.com.

**Figure 2: Structure- and sequence-based protein feature analysis. a**. Enrichments and depletions for protein structural and functional features used by the LoGoFunc model. GOF (blue), LOF (orange), and neutral (green) log odds ratios are displayed for each feature. Significant enrichments and depletions are denoted by asterisks. Significance was calculated with Fisher’s exact test, Benjamini-Hochberg corrected^67^ to allow for multiple comparisons. (Left) Features derived from protein sequences or protein interaction data. (Right) Features derived from AlphaFold2^10^ protein structures. **b**. AlphaFold2^10^ predicted structure of the Vasopressin V2 receptor protein. (Left) Residues colored by the number of HGMD^12^ pathogenic variants occurring in the nine closest neighboring residues in space. (Right) Residues colored by the number of gnomAD^13^ variants occurring in the nine closest neighboring residues in space.

**Figure 3: Association between variant type and impact on splicing. a**. (Top) Density of GOF, LOF and neutral variants within 20 base-pairs of a splice junction. (Bottom) Proportion of GOF, LOF, and neutral variants predicted to yield a gain of splice acceptor or donor or a loss of splice acceptor or donor. **b**. Percentage of GOF, LOF, and Neutral variants in proximity (20 base-pairs) to acceptor and donor splice sites. **c**. MMSplice^21^ sub-model alternate minus reference logit percent-spliced-in predictions for variants predicted to impact splicing. **d**. Distance to the nearest exon junction boundary in nucleotides by variant class. Boxes denote quartiles, whilst whiskers extend to the limits of the distribution with outliers not shown when greater than 1.5 times the interquartile range from the low and high quartiles respectively. Created with BioRender.com.

**Figure 4: Benchmarking LoGoFunc**. Precision-recall curves comparing the discriminatory power of various pathogenicity prediction methods and LoGoFunc on a set of variants from the test set for which predictions were available from all compared tools. **a**. LOF (n. 545) *vs*. neutral (n. 411). **b**. GOF (n. 136) *vs*. neutral (n. 411). **c**. GOF (n. 136) *vs*. LOF (n. 545). **d**. GOF (n. 136) and LOF (n. 545) combined *vs*. neutral (n. 411).

**Figure 5: Explanation of LoGoFunc predictions. a**. SHAP values by class for features with combined SHAP values in the 90^th^ percentile and above. **b**. (Top) The SHAP values for the top ten features for the seven GOF variants found in the ion channel SCN2A in the test set. (Middle) The SHAP values for the top ten features for the eight LOF SCN2A variants in the test set. (Bottom) The SHAP values for the top ten features for the seven neutral SCN2A variants in the test set. **c**. The experimentally determined structure of SCN2A^43^ with the represented GOF (red), LOF (blue), and neutral (yellow) SCN2A variants from the test set. **d**. The SCN2A model from the AlphaFold2 prediction database annotated with the represented GOF (red), LOF (blue), and neutral (yellow) SCN2A variants from the test set.

**Figure 6: Relationship between variant type and phenotypes. a**. Associations between high-confidence, predicted GOF (c.7471C>T) and LOF (c.1648G>A) variants and phenotypes as determined by PheWAS analysis of patients in the BioMe biobank. **b**. Distribution of CADD^4^ PHRED scores in the dataset (green). CADD^4^ PHRED scores and MSC^38^ 95% CI cutoffs for c.7471C>T (solid and dashed blue lines) and c.1648G>A (solid and dashed red lines).

## Notes

### Competing Interest Statement

The authors have declared no competing interest.

### Summary of Updates

Classifier updated to include features derived from AlphaFold2. Additional precalculated predictions for missense variants added.

