## Supplementary Information for "Genome-wide prediction of pathogenic gain- and loss-of-function variants from ensemble learning of a diverse feature set"

#### SUPPLEMENTARY METHODS

#### ANNOTATION

*Conservation*. Primate, mammal, and vertebrate PhastCons[1], and PhyloP[1] scores excluding *Homo sapiens*, were obtained as precalculated annotations provided by CADD v1.6[2] for all human missense mutations in the GRCH38 reference genome and selected indels from gnomAD[3]. PhastCons[1] and PhyloP[1] scores including *Homo sapiens* in the alignments were obtained from dbNSFP[4] via VEP[5]. Grantham[6], Ex[7], PAM250[8], JM[9], and VB[10] substitution scores were obtained from a SQL database of protein annotations provided by SNVBox[11]. dbNSFP[4] was queried via VEP[5] to obtain GERP++[12] and SiPhy_29way[13] conservation scores. PSIC[14] scores for wild-type and mutant amino acids and the number of observed amino acids at the substitution position were obtained as precomputed annotations provided by PolyPhen-2[15]. GERP[16] conservation scores were procured from VEP[5] and the CADD v1.6[2] database. Finally, conservation metrics based on MMSeq2[17] alignments were queried from the DescribePROT[18] database for the human proteome.

*Gene and protein sequence.* Percent GC and CpG in ±75 base-pair windows around the mutation sites were obtained from the CADD v1.6[2] annotation database. cDNA positions, coding sequence positions, protein positions, coding sequence strand and nucleotide position in the codon were obtained with VEP[5]. VEP[5] was also used to determine the distance from the nearest exon junction boundary within 10,000 base-pairs and the length of the nearest exon. Distances to the nearest transcription start sites and transcribed sequence ends were obtained from CADD v1.6[2].

*Genomic motifs, domains, states, and functional regions.* The number of overlapping motifs, highly informative motif positions, and motif score changes when transitioning from wild-type to mutant allele were obtained from the CADD v1.6[2] annotation database. Proximity to splice sites and the nearest mutation in BRAVO were also collected from CADD[2]. Finally, overlapping regulatory features, transcription factor binding sites, chromatin states, and miRNA target predictions were obtained from the CADD v1.6[2] annotation database.

*Allele Frequencies and Mutational Occurrence.* Allele frequencies from the 1000 Genomes Project[19] and the UK10K TWINSUK cohort[20] were collected via the dbNSFP[4] plugin for VEP[5]. The number of frequent (MAF > 0.05), rare (MAF < 0.05), and single occurrence SNVs in BRAVO in 100 and 1000 base-pair windows were obtained from CADD[2]. The frequencies of missense substitution types from COSMIC[21] and HapMap[22] as well as the frequency of missense substitution types in COSMIC[21] normalized by frequency of the reference residue in human proteins in SwissProt/TrEMBL[23] and normalized by the number of times the substitution type was identified in HapMap[22] were queried from SNVBox[11].

*Protein Structure and Composition*. Predictions of protein accessible surface area, disordered flexible linker residues, disordered RNA, DNA, and protein binding residues, RNA and DNA binding residues, MoRF regions, protein secondary structures, protein binding residues, signal peptides, and intrinsically disordered residues, were obtained from the DescribePROT[18] database for the human proteome. Post-translational modifications were collected from iPTMnet[24] and dbPTM[25]. NetSurfP version 1.0d[26] was used to calculate additional estimates of relative and total accessible surface area along with reliability estimates for those scores. We calculated additional predictions of disordered protein regions and disordered binding regions with IUPred2A[27] and ANCHOR2[27], respectively. Pfam[28] and Interpro[29] protein domains were queried from the Ensembl BioMart[30]. Changes in residue side chain volume and solvent accessible surface area, normalized B-factor for the residue, number of hydrogen sidechain-sidechain and sidechain-mainchain bonds formed by the residue, and the average number of residue contacts with heteroatoms per homologous PDB chain along with the closest residue contact with a heteroatom were obtained from the PolyPhen-2[15] annotation database. The average number of residue contacts with other chains per homologous PDB chain, the closest residue contact with another chain, the average number of residue contacts with critical sites per homologous PDB chain, and the closest residue contact with a critical site, were also collected from PolyPhen-2[15]. Documented modified residues and regions of interest were obtained from UniProt[23].

*Genic characterizations*. Predicted and experimentally derived haploinsufficient genes, mode of inheritance predictions, gene selective pressure estimates, genic tolerance to variation, and gene damage metrics were obtained from their respective web servers (Supplementary Table 1). Numbers of paralogs per gene were queried from Ensembl BioMart[30].

*Pathogenicity and deleteriousness.* SIFT4G[31], BayesDel[32], ClinPred[33], DANN[34], DEOGEN2[35], Eigen[36], FATHMM[37], LINSIGHT[38], LIST_S2[39], LRT[40], M_CAP[41], MPC[42], MVP[43], MetaLR[4], MetaSVM[4], MutPred[44], MutationAssessor[45], MutationTaster 2[46], PROVEAN[47], PrimateAI[48], VEST4[49], MSC[50], and fitCons[51] estimates of pathogenicity and deleteriousness were obtained with VEP[5] and the dbNSFP[4] plugin. Variant consequences and their relative impact were also retrieved from VEP[5]. CADD[52], PolyPhen-2[15], SIFT[53], and CONDEL[54] scores were obtained with VEP[5].

*Splicing.* SpliceAI[55] and MMSplice[56] scores were obtained from the CADD v1.6[2] annotation database along with dbscSNV[57] Adaboost and RandomForest classifier scores. MaxEntScan[58] scores were collected via the eponymous VEP[5] plugin.

*Expression and epigenetics.* ENCODE[59] histone modification levels, chromatin state characterizations, and overlapping transcription factor binding sites and regulatory features were obtained from CADD v1.6[2]. Median transcripts per million across 54 tissue types were obtained from GTEx.

*Protein-protein interactions.* The protein-protein interaction network defined across evidence channels from the STRING[60] database version 11 was dimensionally reduced to 64 features with the node2vec[61] implementation found at https://github.com/eliorc/node2vec with default parameters.

#### HYPERPARAMETER TUNING

Hyperparameters were tuned using the Optuna[62] optimization library version 2.10.0 with the Tree-structured Parzen Estimator to sample from the hyperparameter search spaces. We used the LightGBM[63] implementation version 3.2.1 from https://github.com/microsoft/LightGBM, and the scikit-learn[64] API of the XGBoost[65] implementation version 1.5.0 from https://github.com/dmlc/xgboost. We used the RandomForest[66] implementation from scikit-learn[64] library version 1.1.1. Neural networks were implemented using pytorch[67] version 1.8.0. For LightGBM[63], XGBoost[65], and RandomForest[66], parameters in the search spaces below correspond to the parameters defined in the documentations of the implementations located at https://lightgbm.readthedocs.io/en/latest/Parameters.html, https://github.com/dmlc/xgboost/blob/master/doc/parameter.rst, and https://scikit-learn.org/stable/modules/generated/sklearn.ensemble.RandomForestClassifier.html respectively. For the neural network implementations, "n_layers" is the number of hidden layers plus 1, "batchnorm_layerN" determines whether to apply batch normalization to the outputs of the Nth layer, and "dropout_layerN" and "n_units_layerN" refer to the dropout frequency applied to nodes in the Nth layer and the number of nodes in the Nth layer respectively. "activation" is the activation function applied to node outputs, "optimizer" is the optimizer employed by the model, "lr" is the learning rate, and "weight_decay" is the L2 regularization penalty.

*LightGBM[63] search space.*

num_iterations: Integer between 100 and 1000. Step size of 50.
num_leaves: Integer between 2 and 3002. Step size of 20.
learning_rate: Floating point number between .01 and .3.
min_child_weight: Floating point number between .01 and 20.
min_data_in_leaf: Integer between 5 and 100.
max_depth: Integer between -1 and 20.
colsample_bytree: Floating point number .4 and 1.
Subsample: Floating point number .4 and 1.
reg_lambda: Integer between 0 and 100.
reg_alpha: Integer point number between 0 and 100.
is_unbalance: Boolean.

*RandomForest[66] search space.*

n_estimators: Integer between 100 and 1000. Step size of 100.
max_features: One of: "auto", "sqrt", "log2".
max_depth: Integer between 1 and 200 or "None".
min_samples_split: Floating point number between .00001 and 1.
min_samples_leaf: Integer between 1 and 20.
booststrap: Boolean.

*XGBoost[65] search space.*

learning_rate: Floating point number between .001 and .401. Step size of .001.
min_child_weight: Integer between 0 and 200.
max_depth: Integer between 1 and 200.
colsample_bytree: Floating point number between .4 and 1. Step size of .1 .
colsample_bylevel: Floating point number between .4 and 1. Step size of .1 .
max_delta_step: Integer between 0 and 10.
gamma: Floating point number between 0 and 100.
reg_lambda: Integer between 0 and 100.
reg_alpha: Integer between 0 and 100.
subsample: Floating point number between .4 and 1. Step size of .1.

*Neural Network search space.*

n_layers: Integer between 2 and 5.
batchnorm_layerN: One of: “True”, “False”.
n_units_layerN: Integer between 128 and 2,056.
dropout_layerN: Floating point number between 0 and .9. Step size of .01.
activation: One of: “LeakyReLU”, “ReLU”, “Sigmoid”, “Tanh”.
optimizer: One of: “Adam”, “RMSprop”, “SGD”.
lr: Floating point number between .00001 and 1 sampled from the log-uniform distribution.
weight_decay: One of: .00001, .0001, .001, .01, .1, 0.
batch_size: Integer between 256 and 1,024.
do_classweight: Boolean indicating whether to weight samples inversely proportional to class frequencies.

### SUPPLEMENTARY FIGURES

#### SUPPLEMENTARY FIGURE 1

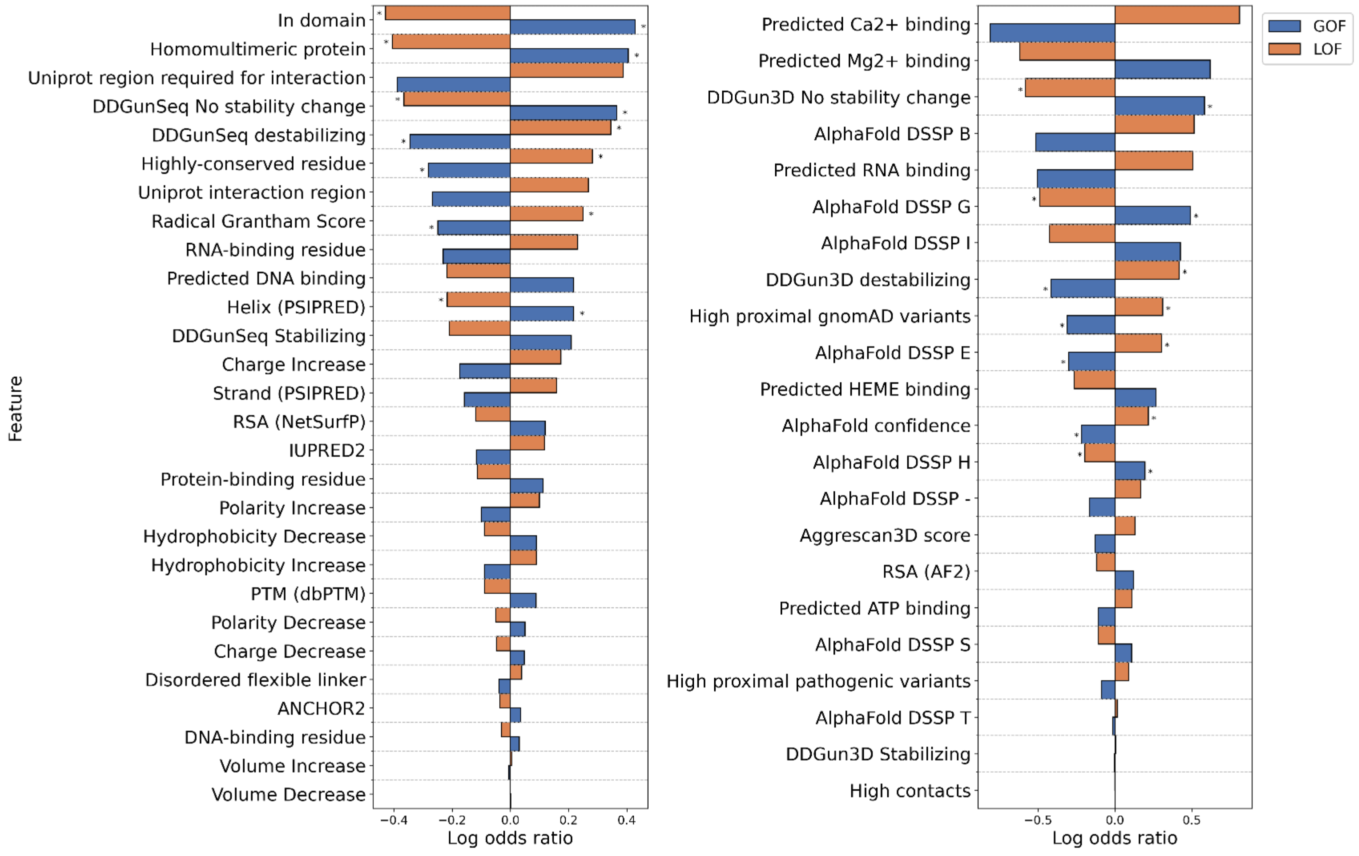


**Supplementary Figure 1**: Enrichments and depletions for protein structural and functional features used by the LoGoFunc model. GOF (blue) and LOF (orange) log odds ratios are displayed for each feature. Significant enrichments and depletions are denoted by asterisks. Significance was calculated with Fisher's exact test, Benjamini-Hochberg[68] corrected to allow for multiple comparisons. (Left) Features derived from protein sequences or protein interaction data. (Right) Features derived from AlphaFold2[69] protein structures.

#### SUPPLEMENTARY FIGURE 2

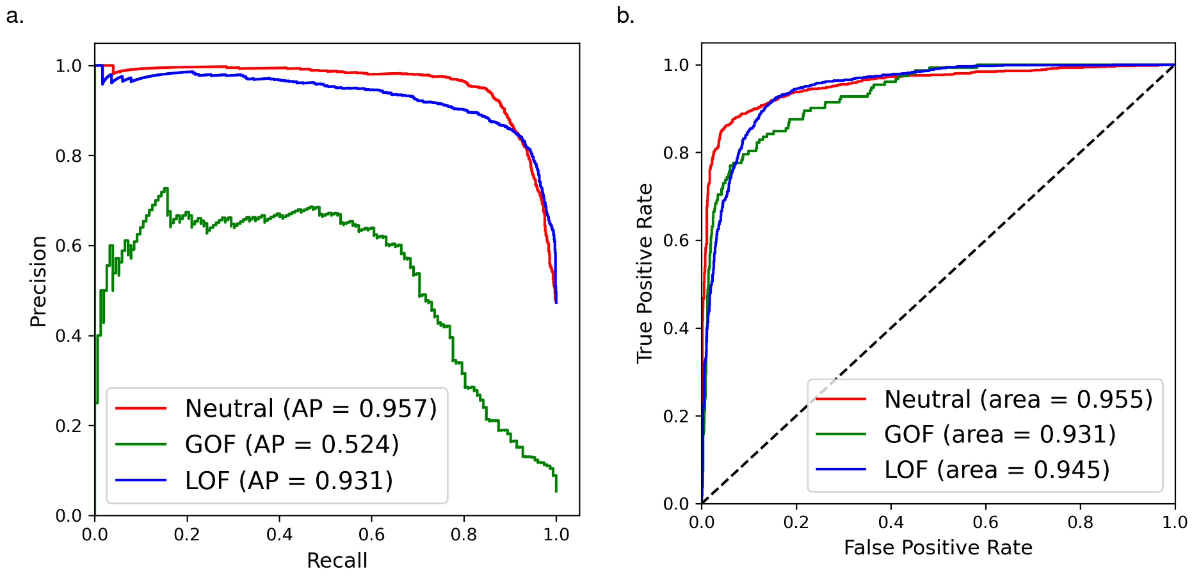


**Supplementary Figure 2**: **a.** Precision-recall curves by variant class for all variants in the test set. **b.** Receiver operating characteristic curves by variant class for all variants in the test set.

#### SUPPLEMENTARY FIGURE 3

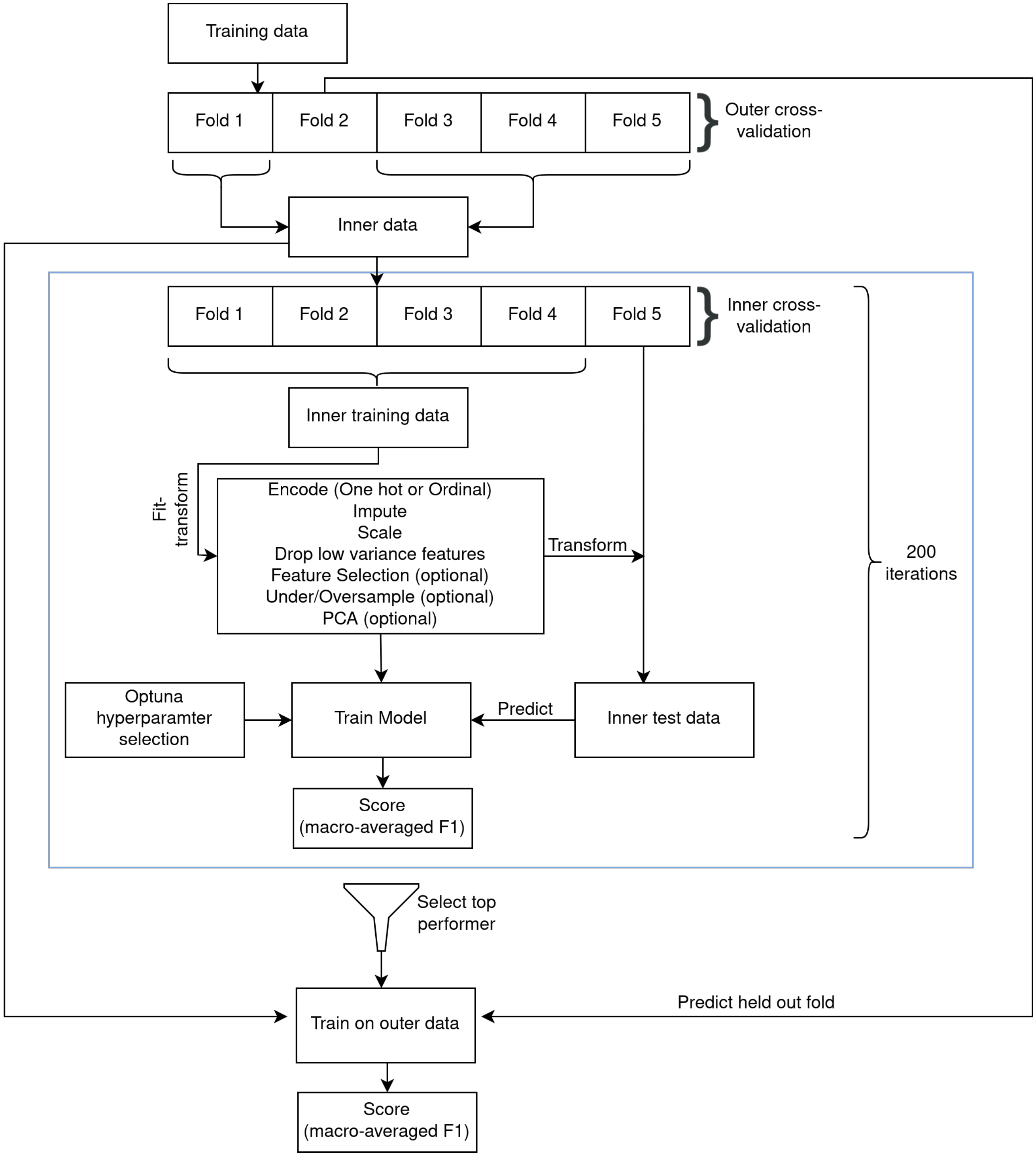


**Supplementary Figure 3**: Nested cross-validation strategy employed for model selection. Illustration of the nested cross-validation strategy employed for model selection in which fold two is held out for an iteration of the outer validation loop and fold five is held out for an iteration of the inner loop. Folds are stratified by variant class and split such that the set of genes in each fold is disjoint. Preprocessing steps and hyperparameters are tuned in the inner loop and performance is assessed by the macro-averaged F1 score in both the inner and outer loops.

#### SUPPLEMENTARY FIGURE 4

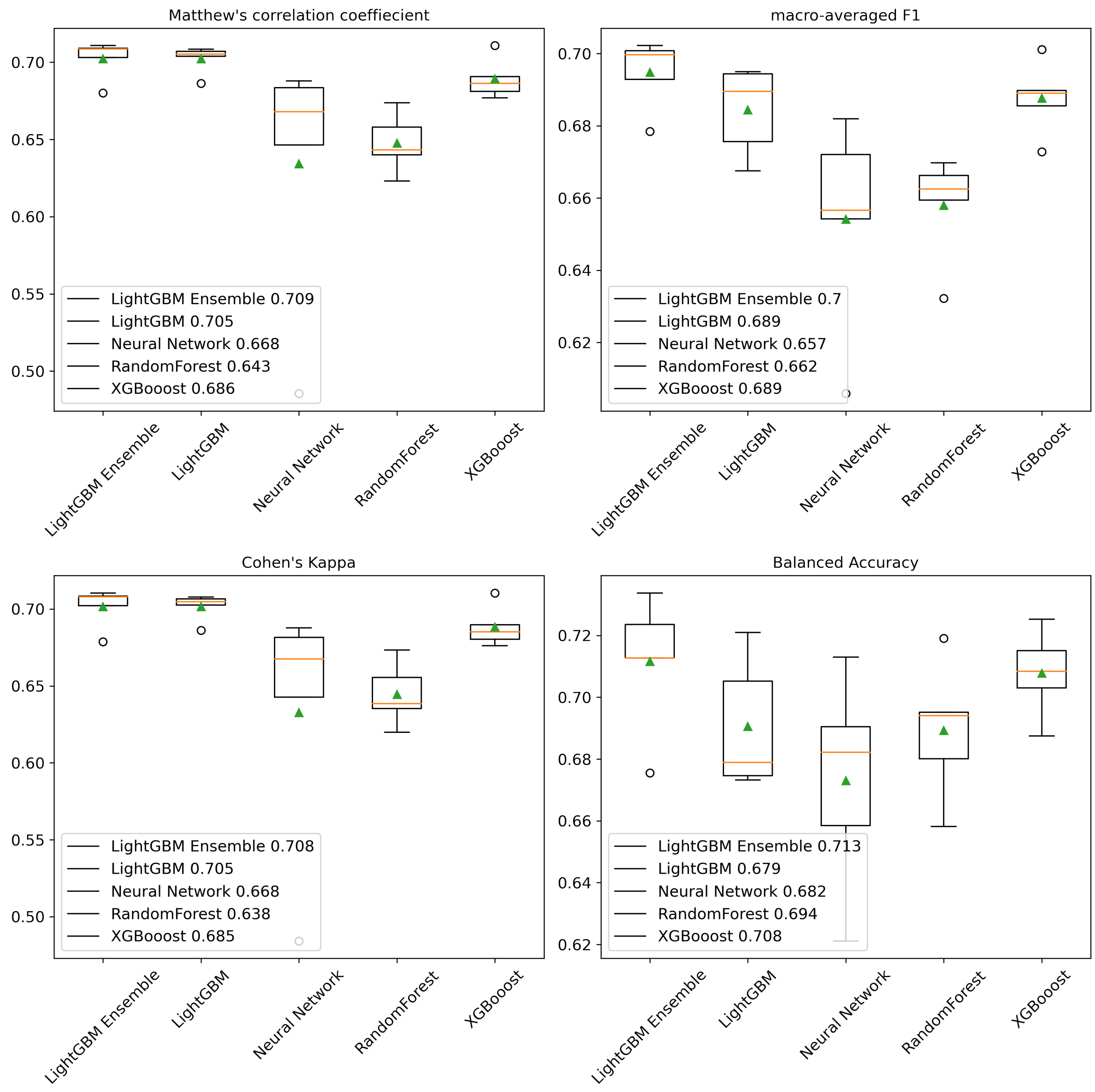


**Supplementary Figure 4**: Box and whisker plots for the comparison of tested model architecture performance on the outer folds of the nested cross-validation loop. Models are compared by respective Matthew's Correlation Coefficient, Cohen's kappa, macro-averaged F1, and balanced accuracy scores. Green triangles denote the mean score by model and orange horizontal lines denote the median score by model. Boxes extend from the first to the third quartiles. Whiskers extend by 1.5 times the interquartile range past the boxes. Values beyond the whisker range are denoted by circles.
